# Sharing power: effects of rider ability and position on tandem performance

**DOI:** 10.64898/2026.03.25.714296

**Authors:** Albert Smit, Jip van Ewijk, Ina Janssen, Thomas W.J. Janssen, Mathijs J. Hofmijster

**Author notes:** Corresponding author, (AS).

## Abstract

**Objective:** Tandem cycling requires a coordinated effort between the pilot and the stoker. Previous research suggests that randomly paired tandem cyclists produce lower power output than when cycling solo. This study examined how a cyclist’s individual ability and their position on the tandem (pilot or stoker) affects pair performance, when partners are either closely matched or differ substantially in solo cycling capacity, as this might be relevant for training and selection.

**Methods:** Twenty-three trained cyclists completed three 10-minute time trials: solo, equal-capacity tandem (≤25 W difference in solo performance), and unequal-capacity tandem (≥40 W difference). Mean power output, heart rate, cadence, and rating of perceived exertion (RPE) were recorded. Positions (pilot or stoker) were counterbalanced. Linear mixed-effects models assessed effects of capacity and position.

**Results:** Relative to solo cycling, equal-capacity tandem pairs revealed lower power output (-3.9%), lower heart rate (-2.3%), and lower RPE (-11.5%). Unequal-capacity tandems differed from solo only in heart rate (-2.7%). Stokers produced lower power relative to solo (-5.3%) and relative to pilots (-3.7%) and reported lower RPE relative to solo (-13.9%), while pilots matched their solo power at a lower heart rate (-2.9%). Cadence did not differ across conditions. Total tandem power averaged 95.6% of combined solo power, and differences in partner capacity did not significantly affect combined power output.

**Conclusion:** This study provides the first known experimental data on how partner matching affects individual and combined power output in tandem cycling. Equal- and unequal-capacity tandem pairs showed similar performance. Lower power and RPE among stokers suggest reduced engagement or a redistribution of effort between riders. These findings highlight that effective tandem performance depends on physiological capacity and rider position on the tandem, but not on the difference in capacity between partners.

## Introduction

Tandem cycling is a Paralympic discipline in which a cyclist with a visual impairment (the stoker, seated at the rear) rides with a sighted pilot (seated at the front), who controls steering, braking, and gear shifting. Both pilot and stoker contribute to the total power output. Ideally, total tandem power approaches the sum of the of the individual cyclists’ net mechanical power output (PO), although this is rarely achieved [1]. Despite its competitive relevance, the physiological and mechanical determinants of optimal tandem performance remain poorly understood [2].

In a previous study, we compared solo and tandem cycling performance using randomly assigned partners [1]. We observed decreased PO and increased cadence during tandem cycling relative to solo cycling efforts. Large within-pair differences in maximal PO raised the question of whether pairing itself affected performance. However, that dataset lacked sufficient statistical power to assess this relationship.

We hypothesized that the degree of partner matching influences PO and overall tandem performance. Large discrepancies in performance capacity may reduce efficiency as a pair, as shown in tandem cycling research with Parkinson’s patients, where stoker passivity increased power dissipation [3]. In addition to this, Seifert et al. [4] reported that during submaximal efforts, higher capacity cyclists maintained similar physiological responses across solo and tandem rides, whereas lower capacity cyclists displayed lower heart rate and perceived exertion. These effects were most pronounced among stokers, consistent with the possibility that both partner mismatch and position contribute to performance differences.

Pacing, the goal-directed regulation of exercise intensity across an effort [5], is a key determinant of endurance performance [6]. However, as riding with a partner on a tandem alters PO [1], it is likewise plausible that a partner also influences pacing behaviour on a tandem, as it does in for example sprint kayak, where kayaks with 4 persons show a different pacing profile compared to single-person kayaks [7]. So far, the influence of tandem cycling on pacing remains unexplored.

In the present study we (1) aimed to evaluate whether differences in solo cycling capacity between partners influence individual and combined tandem performance, and (2) to examine how rider position (pilot vs. stoker) affects physiological and perceived exertion. As a third objective, we examined the influence of tandem cycling and tandem partners on pacing strategies. Specifically, we investigated the effects of partner matching (equal-vs. unequal-capacity), hypothesizing that pairs with equal-capacity would perform relatively better on a tandem than partners with unequal-capacity. To this end, we assessed PO, cadence, heart rate, ratings of perceived exertion (RPE), and pacing strategy during both solo and tandem trials. We also investigated the effects of relative capacity within unequal-capacity pairs (higher vs. lower capacity), and the influence of position (pilot vs. stoker) on the same parameters. This study addresses a key gap in understanding how pairing strategies affect combined PO production in tandem cycling. These findings may assist coaches and athletes in optimising partner selection and training strategies to enhance tandem cycling performance.

## Methods

### Study Design and Participants

We employed a within-subject, controlled crossover design in which each participant completed cycling trials under three conditions across separate sessions: (1) solo, (2) tandem with a partner equally matched on solo PO, and (3) tandem with an unequally matched partner on solo PO. For the tandem condition, each participant was assigned to the stoker or pilot position for sessions 2 and 3 in a counterbalanced fashion. The target sample size (n ≥21) was based on an a-priori power analysis indicating that, with power >0.80 and α <0.05, this sample size was sufficient to detect medium-sized effects (d = 0.6). The expected effect size was based on the magnitude of the solo-tandem differences in our previous study [1].

Twenty-three trained cyclists (1 female, 22 males), none of whom had prior tandem experience, participated in the study and were recruited between 18-03-2025 and 01-05-2025. All participants executed the solo condition first to obtain individual capacity, expressed as average PO over the complete time trial. Trial order of the tandem conditions was randomised and counterbalanced across participants to minimise order effects, All participants completed a Physical Activity Readiness Questionnaire (PAR-Q) [8] to screen for contraindications to maximal exercise and provided written informed consent after a detailed briefing. This study was approved by the Ethics Committee of the Faculty of Behavioural and Movement Sciences at Vrije Universiteit Amsterdam (VCWE-2025-068) and was conducted in accordance with the Declaration of Helsinki.

### Experimental design

Each participant completed three sessions, separated by at least 24 hours (and typically ≥48 hours) to minimise fatigue. Participants were instructed to avoid vigorous exercise for 24 hours before testing. During the first session, body mass and height were measured using a Seca robusta 813 scale (Seca, Germany) and HM-250P stadiometer (Marsden, UK). Then, after a standardised warm-up, each cyclist completed a 10-minute solo time trial on a tandem bicycle mounted on a Cyclus2 ergometer (RBM electronic-automation GmbH, Leipzig, Germany), which had a direct-current motor replacing the rear wheel. Saddle height (centre of bottom bracket to top saddle, inline with the saddle tube) was recorded to the nearest millimetre for replication. The participants wore their own cycling shoes with SPD-SL cleats. PO, heart rate, and cadence were recorded at 1 Hz, and RPE was obtained every two minutes on a 11-point visual scale [9].

In subsequent sessions, participants were paired based on mean solo PO. Participants were assigned to an equal-capacity partner (equal-capacity tandem), in which partners differed by ≤25 W, and to an unequal-capacity partner (unequal-capacity tandem), in which partners differed by ≥40 W. These thresholds were selected to create clearly distinguishable pairing conditions while allowing sufficient pair formation within the available sample. In the unequal-capacity tandem condition, each individual was either the higher-capacity or the lower-capacity cyclist. Each participant completed a 10-minute tandem time trial in both pairing conditions, performing one as pilot and one as stoker. Pilot and stoker positions were equally distributed across higher- and lower-capacity cyclists, meaning that during the unequal-capacity tandem trials, 50% of the pilots were higher-capacity and 50% of the pilots were lower-capacity cyclists.

### Equipment and Data Collection

All trials were conducted using the same road tandem bicycle (Duratec, s.r.o., Czech Republic) mounted on the same ergometer. PO was obtained using Garmin Rally RS200 dual-sided pedals (Garmin International, Inc., Olathe, Kan, USA), which have a typical measurement error of 3–5% for power and 3% for cadence [10, 11]. Heart rate was monitored using Garmin HRM-Dual straps, and all data (power, cadence, heart rate) were recorded using Garmin Edge 530 devices. Pedals were offset-corrected before each trial according to manufacturer instructions.

During solo trials, cyclists rode the tandem at the position of the pilot and controlled cadence and gear shifting. No specific pacing or cadence instructions were given. In the tandem conditions, participants were encouraged to communicate before and during trials to coordinate their effort, allowing both pilot and stoker to influence gear selection, with the pilot operating gear shifting. Tests were conducted in a thermoneutral environment, and a fan was used to stimulate airflow across the participants.

### Test Protocols and Measurements

Each session started with a standardised warm-up consisting of four 3-minute stages, starting at 100 W individual PO, and increasing with 25 W each stage, at 90–100 rpm, followed by 3 minutes of active recovery at 75 W. For the tandem trials, the imposed external load on the tandem pair was doubled to simulate an equivalent load per rider, and participants were instructed to share the workload evenly by applying similar PO. A 2-minute familiarisation period followed the warming-up, allowing the pair to select the gear to start with.

The aim of the time trial was to achieve the highest mean PO possible in 10 minutes. All time trials were conducted in “inclination mode”, which simulated outdoor cycling dynamics, including air resistance, rolling resistance, gravity, and inertia. Real-time feedback on cadence, heart rate, PO, and elapsed time was provided to each participant via the display of the cycle computer. Verbal reminders of the remaining time were given every two minutes, with additional cues during the last minute (“One minute to go”; “Thirty seconds to go”; “Fifteen seconds to go”). RPE (0–10 scale; [9] ) was recorded every two minutes following the standard verbal prompts.

### Statistical analysis

Linear mixed-effects models were fitted to account for the repeated-measures design, with participant ID as the random intercept to account for non-independence of observations. To ensure stable estimation and avoid overfitting, simplified random structures (no random slopes) were used after verifying model convergence. Normality and homoscedasticity were verified visually using Q-Q plots and residual-vs-fitted plots. All models were fitted using restricted maximum likelihood.

PO and heart rate were normalised to individual solo performance by dividing the tandem values by each cyclist’s mean solo output. The effects of riding configuration (solo, equal-capacity tandem, unequal-capacity tandem, higher-capacity, lower-capacity) and rider position (pilot, stoker) on relative power, relative heart rate, mean RPE, and cadence (equal for pilot and stoker and therefore analysed for riding configuration only) were evaluated in separate models. Bonferroni-adjusted pairwise comparisons were used to obtain post-hoc contrasts.

To assess pacing, RPE and 2-minutes mean power (0–2, 2–4, 4–6, 6–8, 8–10 minutes) were modelled separately with time and riding configuration and time and position and their interaction (time × configuration or time × position) as fixed effects. Random slopes for time were excluded due to singular model fits.

To evaluate how differences in individual performance capacity influenced overall pair PO, we calculated combined relative performance for each tandem pair as the total mean PO on the tandem divided by the sum of both riders’ solo mean PO. This analysis quantified how effectively pairs performed as a unit. We then assessed whether between-cyclist differences in solo power were related to combined performance.

Because PO was sampled continuously over time, adjacent observations were temporally dependent; therefore, cluster-based permutation tests were used to account for temporal dependency in the time series [12]. As pilot and stoker positions were not repeated within participants within a given condition, differences in relative PO between positions were assessed using cluster-based permutation tests for independent samples. These analyses were performed separately for equal-capacity tandems and for higher- and lower-capacity cyclists in the unequal-capacity tandem condition.

Effect sizes (Cohen’s *d*) were computed as the estimated difference divided by the residual Standard Deviation (SD) [13]. Omega squared (ω^2^) was reported for main effects as a less biased measure for small samples [14, 15]. All analyses were conducted in JASP (v0.95.3.0; JASP Team, Amsterdam, The Netherlands) and Python (v3.12; Python Software Foundation) using standard available scientific libraries. Probability level for statistical significance was set at p< 0.05 (two-tailed). All data will be deposited in an open repository at publication, in accordance with the PLOS ONE Data Policy.

## Results

All participants were classified as Tier 2 (“Trained”) according to established performance guidelines [16] (Table 1). A total of 24 equal-capacity tandem and 18 unequal-capacity tandem trials were completed, as not all participants could be unequally matched due to too small differences in PO. Heart rate data were incomplete for three cyclists and were excluded from analyses involving this variable. Subsequent analyses examined PO, heart rate, RPE, and pacing across riding configurations and positions. Descriptive statistics for power output, cadence, heart rate, and perceived exertion across riding configurations and rider positions are presented in Table 1, while estimated marginal means from the linear mixed models for power and heart rate relative to solo and for RPE are shown in Table 2. Fig 1 illustrates relative PO across configurations and positions. Cadence did not differ significantly across riding configurations (F(2, 42.11) = 1.78, p = 0.182, ω^2^ = 0.03).

**Table 1.** Participant characteristics (mean ± SD)

|  |  |
| --- | --- |
| N | 23 |
| Age (yrs) | 38 ± 17 |
| Training years | 14 ± 10 |
| Training frequency (times per week) | 4.0 ± 2.3 |
| Training duration (hours per week) | 7.3 ± 3.0 |
| Height (m) | 1.84 ± 0.09 |
| Body mass (kg) | 78.3 ± 10.7 |

**Table 1.**
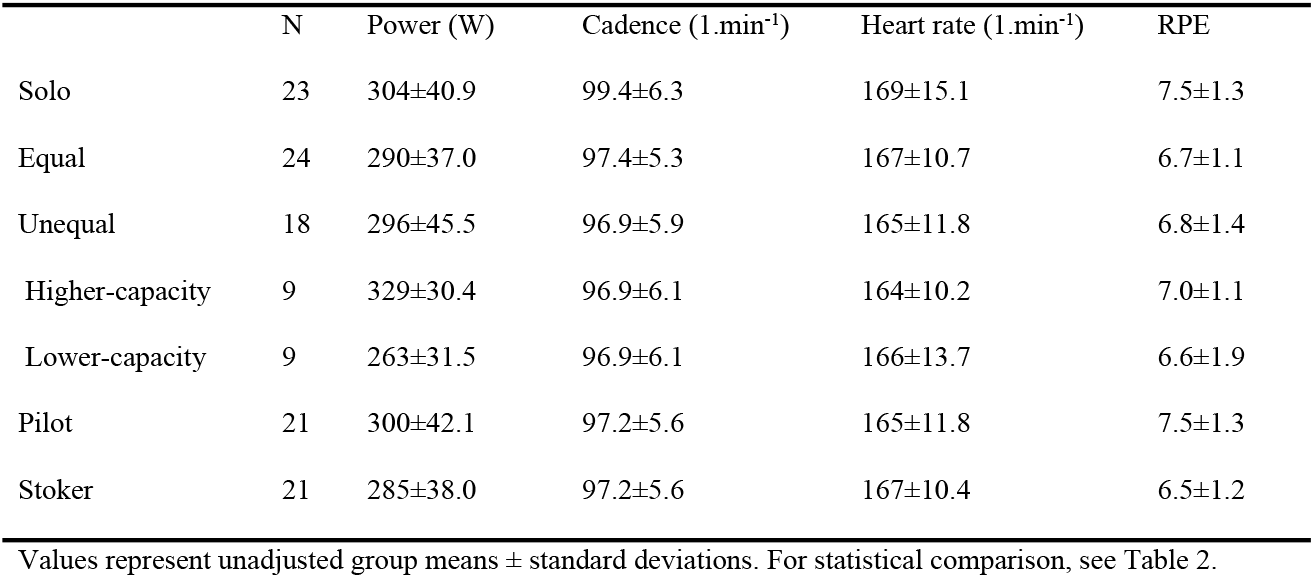
Descriptive statistics for power output, cadence, heart rate, and perceived exertion across riding configurations and rider positions.

|  | N | Power (W) | Cadence (1.min<sup>-1</sup>) | Heart rate (1.min<sup>-1</sup>) | RPE |
| --- | --- | --- | --- | --- | --- |
| Solo | 23 | 304±40.9 | 99.4±6.3 | 169±15.1 | 7.5±1.3 |
| Equal | 24 | 290±37.0 | 97.4±5.3 | 167±10.7 | 6.7±1.1 |
| Unequal | 18 | 296±45.5 | 96.9±5.9 | 165±11.8 | 6.8±1.4 |
| Higher-capacity | 9 | 329±30.4 | 96.9±6.1 | 164±10.2 | 7.0±1.1 |
| Lower-capacity | 9 | 263±31.5 | 96.9±6.1 | 166±13.7 | 6.6±1.9 |
| Pilot | 21 | 300±42.1 | 97.2±5.6 | 165±11.8 | 7.5±1.3 |
| Stoker | 21 | 285±38.0 | 97.2±5.6 | 167±10.4 | 6.5±1.2 |

Values represent unadjusted group means ± standard deviations. For statistical comparison, see Table 2.

**Table 2.** Estimated marginal means for riding configuration and position for power and heart rate relative to solo, and for RPE.

|  | Relative power | Relative heart rate | RPE |
| --- | --- | --- | --- |

|  | [95% CI] | [95% CI] | [95% CI] |
| --- | --- | --- | --- |
| Equal-capacity tandem | 0.96 [0.94 0.98] * | 0.98 [0.96 0.99] * | 6.6 [6.1 7.1] * |
| Unequal-capacity tandem | 0.97 [0.95 0.99] | 0.97 [0.96 0.99] | 6.8 [6.2 7.4] * |
| Higher-capacity | 0.97 [0.94 1.00] | 0.98 [0.95 0.99] | 6.9 [6.2 7.8] |
| Lower-capacity | 0.97 [0.95 1.00] | 0.97 [0.95 0.99] | 6.6 [5.9 7.4] |
| Pilot | 0.98 [0.97 1.00] | 0.97 [0.96 0.99]* | 6.9 [6.4 7.4] * |
| Stoker | 0.95 [0.93 0.97] *# | 0.98 [0.97 1.00] | 6.4 [5.9 7.0] * |

\* Significant difference with solo, $p < 0.05$

**Fig 1.**
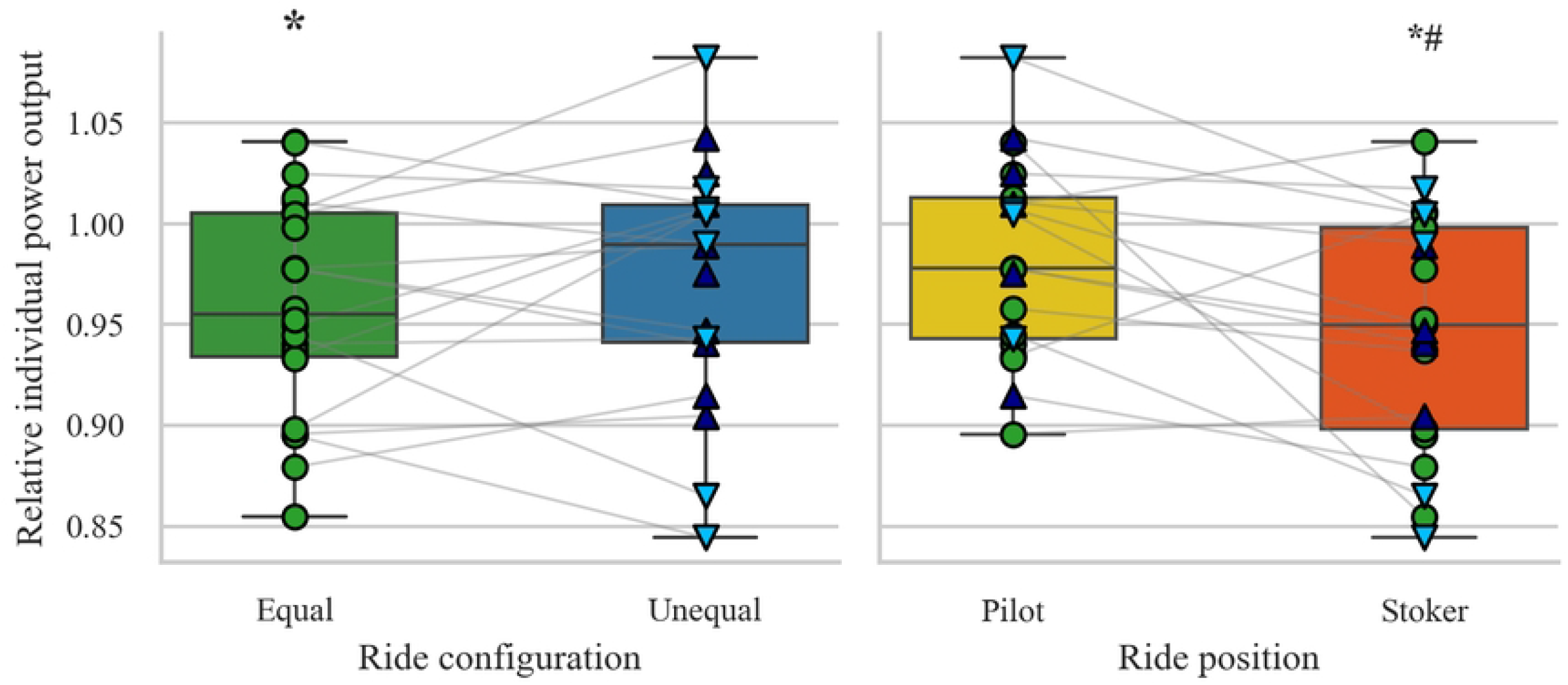
Power output relative to solo performance across conditions and positions. Box plots portray relative power output for ride configuration (equal/unequal; left figure) and ride position (pilot/stoker, right figure). Green circles indicate equal ride, dark blue triangles indicate higher-capacity, and light blue triangles indicate lower-capacity participants. *Indicates significant difference from solo (p <0.01), # indicates significant difference from pilot (p <0.01)

### Relative individual power output

Pairwise contrasts revealed that PO in equal-capacity tandem rides differed significantly from PO in solo rides (-3.9% ≈ 14 W, p = 0.003). No significant differences were observed between unequal-capacity tandem and solo rides, or between equal-capacity tandem and unequal-capacity tandem conditions. Within unequal-capacity tandem pairs, neither higher-capacity nor lower-capacity cyclists differed significantly from their solo performances, nor between the two partners. Regarding rider position, cyclists in the stoker position produced significantly less PO than during solo rides (-5.3% ≈ 19 W, p < 0.001) and less than cyclists in the pilot position (-3.7% ≈ 15 W, p = 0.007), whereas pilot PO did not differ from solo.

### Relative individual heart rate

Individual heart rate was significantly lower during equal-capacity tandem and unequal-capacity tandem rides compared to solo cycling (-2.3% ≈ 2 bpm, p = 0.049 and -2.7% ≈ 4 bpm, p = 0.028, respectively). No differences were found between the tandem conditions, suggesting that all tandem conditions were performed at a lower physiological load than during solo. Post-hoc analysis revealed that pilots exhibited lower heart rates relative to their solo rides (-2.9% ≈ 4 bpm, p = 0.005), while stokers did not differ significantly from their solo rides or from pilots. Thus, while pilot PO was comparable to solo levels, heart rate data suggest that their cardiovascular strain was lower when riding tandem.

### Ratings of perceived exertion (RPE)

Riding configuration significantly affected mean RPE. Equal-capacity tandem rides elicited lower RPE than solo (-11.5%, p = 0.011), indicating that well-matched partners perceived the effort as less demanding. No significant differences were observed between all other tandem configurations. Statistic analysis showed an effect of position on RPE, with stokers reporting significantly lower RPE during tandem cycling than during their solo rides (-13.9%, p = 0.002), with no difference to pilots. This indicates that stokers perceive the effort as less demanding than solo cycling, but comparable to pilots.

### Pacing

A significant main effect of riding configuration (F(2, 288.89) = 7.39, p < 0.001) and time (F(4, 287.93) = 28.01, p < 0.001) was found for PO. No configuration × time interaction was observed (p = 0.382). Mean power in the equal-capacity tandem condition was lower than solo during the first two minutes (-6.5%, p = 0.009). PO was comparable across time in all conditions from 2 minutes onward, with similar pacing patterns toward the end of the trial (Fig 2).

**Fig 2.**
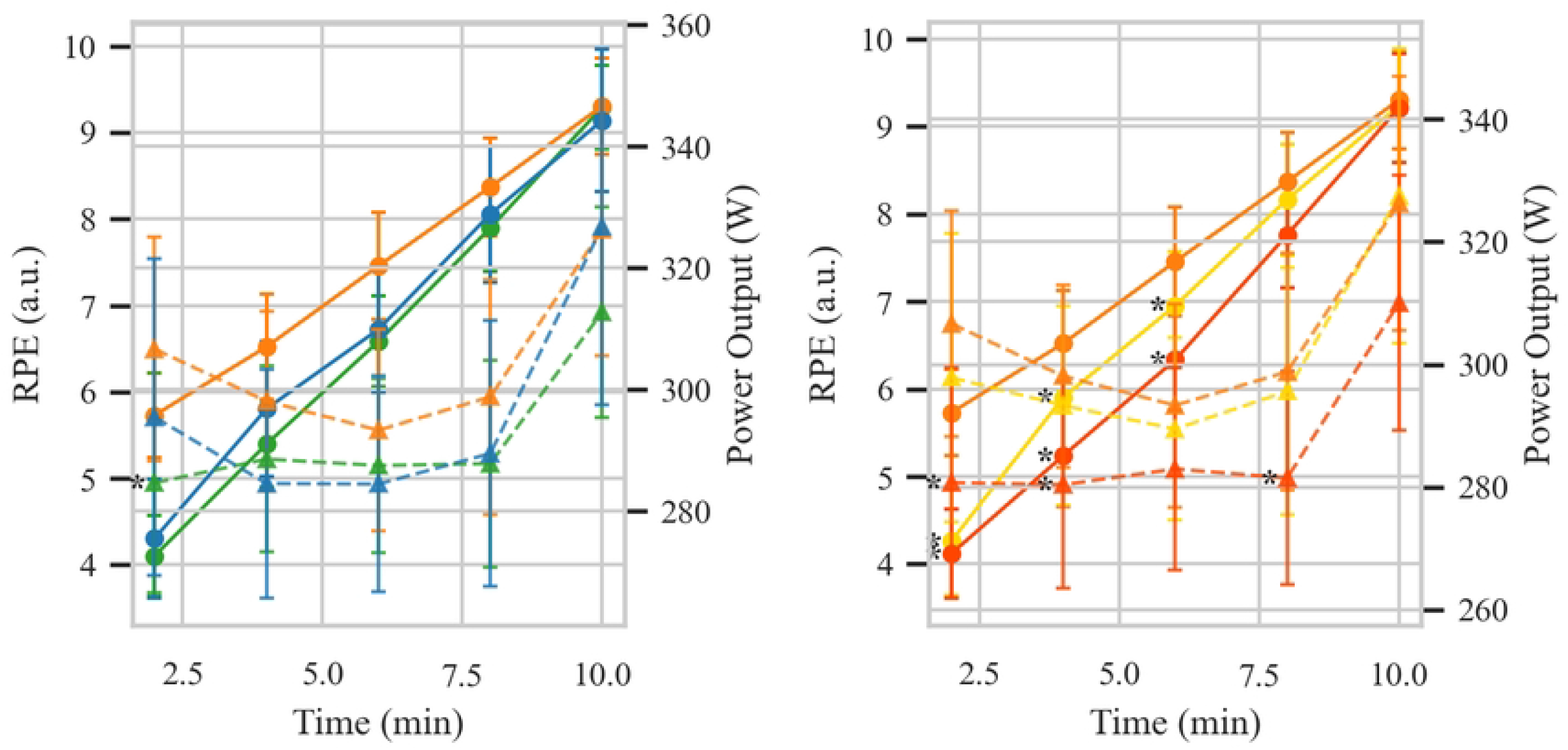
RPE and power output in 2-minute time bins, for riding configuration (left figure) and position (right figure). Solid lines and dots indicate ratings of perceived exertion (RPE), and dashed lines and triangles indicate power output. Orange is solo, green is equal-capacity tandem, blue is unequal-capacity tandem, yellow is pilot and red is stoker. Error bars indicate 95% confidence interval. * indicates significant difference with solo (p < 0.05).

When examining rider position, main effects of position (F(2, 289.18) = 10.74, p < 0.001) and time (F(4, 287.93) = 27.32, p < 0.001) were found; position × time did not interact (p = 0.836). Across time points, stokers demonstrated lower PO than during solo condition (-7.1%, p = 0.030 from 0-2 minutes), while pilots did not differ significantly from solo or stokers. These results indicate that stokers consistently produced lower PO across time, while pacing patterns remained similar across positions.

In the unequal-capacity tandem conditions, pilots produced higher or equal PO compared to solo cycling in the first minute of the time trial (see Fig. S1 A, B, C in the supplement). This was more pronounced when a lower-capacity pilot rode with a higher-capacity stoker, with significant differences between lower-capacity pilots and lower-capacity stokers in the first 13 s (p=0.019). See Fig. S1 C. However, in equal-capacity tandems, the pilots appear to increase PO gradually during the first 120 s, while the stokers keep a pacing strategy comparable to solo cycling from the start (Fig. S1 A).

For RPE, both time (F(4, 287.76) = 174.70, p < 0.001) and riding configuration (F(2, 291.95) = 20.83, p < 0.001) showed significant effects, with a small but significant time × configuration interaction (F(8, 287.76) = 2.16, p = 0.031) (Fig 2). RPE increased from approximately 4 to 9 across trials, except for solo cycling, which started at a higher value (∼6). Solo condition consistently elicited higher RPE than tandem rides between 2–6 minutes (all p < 0.05). By 8–10 minutes, RPE converged across conditions to 9-10. Overall, these results indicate that objective pacing patterns, i.e. expressed as PO, were comparable across configurations, however, subjective effort was consistently lower in tandem conditions, especially in the first half of the time trial.

### Combined relative performance

On average, pairs produced about 95.6% of their combined solo power when riding tandem (mean combined relative performance = 0.956, Standard Error = 0.007). We found no significant relationship between differences in solo capacity and combined relative performance (F(1, 24.87) = 3.56, p = 0.071). This suggests that while individual cyclists may contribute unequally to the joint effort, the summed power of the pair remains largely unaffected by how well matched they are. Thus, mismatched performance was comparable to matched performance, with only minimal loss in overall output (Fig 3).

**Fig 3.**
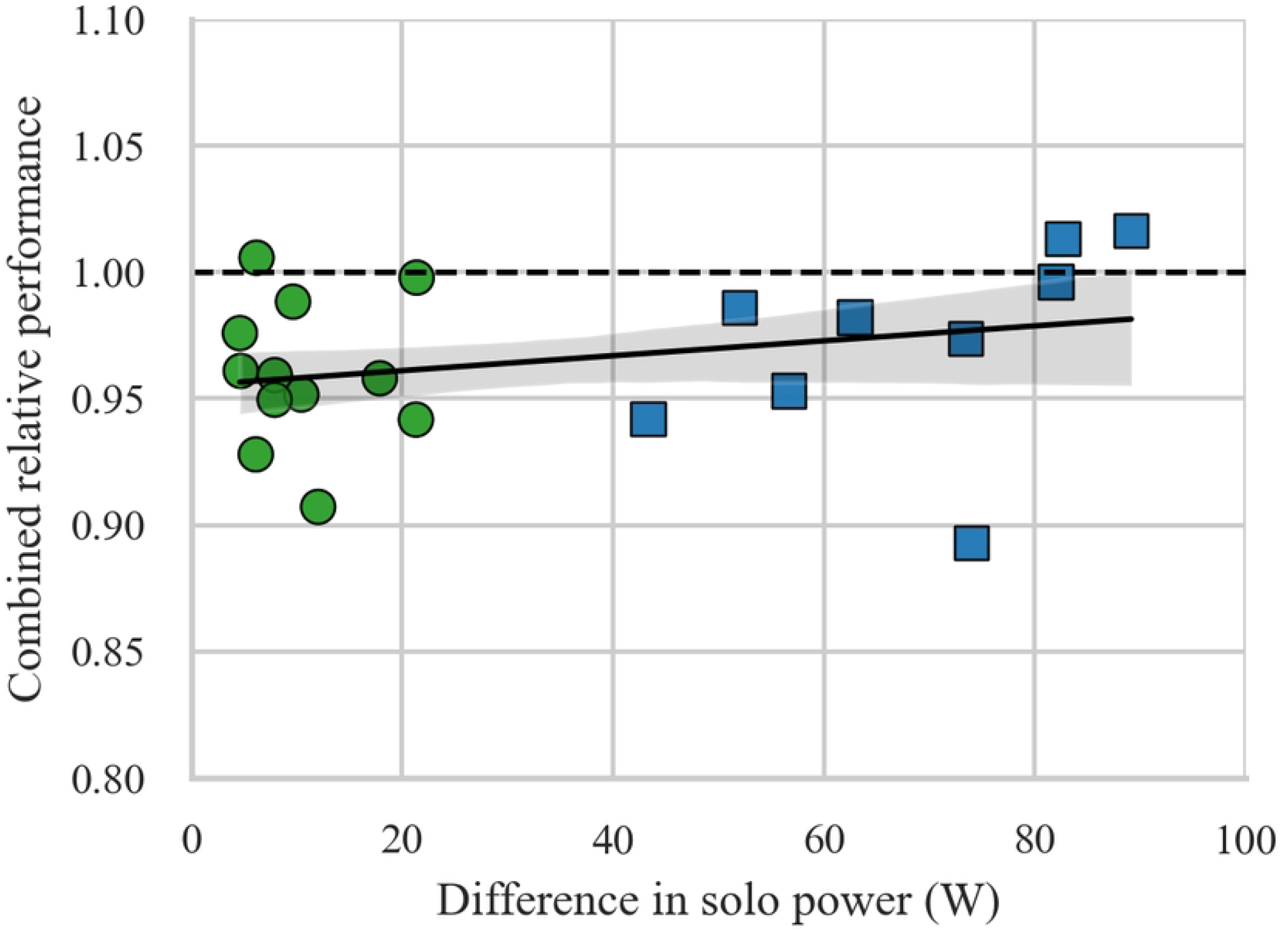
Relation between combined relative performance and difference in solo power. Each data point represents one pair, green circles are equal-capacity tandem pairs, blue squares are unequal-capacity tandem pairs. The black line indicates the regression line and the shaded area the 95% confidence interval for this regression line. The horizontal dashed line indicates combined solo efforts to be equal to tandem effort.

## Discussion

The present study examined how individual performance capacity and rider position influenced tandem time-trial performance in trained but tandem-inexperienced cyclists. Consistent with our previous work [1], PO on the tandem was lower than during solo cycling, particularly in equally matched pairs and among stokers. Together, these findings indicate that tandem performance is influenced not only by individual capacity but also by how effort is regulated between riders.

### Interpersonal dynamics and partner matching

Contrary to our hypothesis, tandem pairs with unequal solo capacities performed no worse than equally matched pairs. On average, pairs produced 95.6% of their combined solo power, and partner mismatch did not significantly predict total output. Interestingly, lower-capacity pilots in unequal-capacity pairs appeared to compensate partially for being the lower-capacity cyclist by starting much harder than during their solo effort, while lower-capacity stokers seemed to underperform at the start of the time trial. This would be consistent with adaptive workload distribution rather than an increase or loss of effectiveness [17]. Similar compensatory mechanisms have been observed in cooperative tasks [18] and group cycling [19], where individuals adjust effort to maintain group cohesion and performance stability. These findings indicate that within the tested range in mismatch (≤25 W vs. ≥40 W difference), differences in individual capacity did not compromise short-duration tandem performance. Instead, the results suggest that riders spontaneously regulate their relative contribution to sustain overall pair output, which can be seen as a form of implicit team effort rather than the simple summation of individual abilities.

At the same time, the lower output observed in equal-capacity tandems and among stokers points to additional influences beyond physiological capacity. While the pilot controlled the gear lever, the explicit task instructions for both pilot and stoker were identical, namely to produce maximal effort. However, with shared responsibility, individual accountability may decrease, leading to under-contribution, a phenomenon consistent with the Ringelmann effect [20-22]. This social dynamic may have been more apparent in the stokers, and could also plausibly arise in equal-capacity tandem conditions where neither rider’s dominance nor dependency is clearly established. Conversely, pilots may benefit from a social facilitation effect, with heightened arousal and engagement through their leadership and control roles [23, 24]. Together, these findings indicate that tandem performance is shaped not only by the physiological compatibility of partners but also by social and motivational dynamics that regulate shared effort.

Unlike in our previous study [1] where we also compared solo and tandem cycling performance, we found in the current study no significant difference in cadence between solo and tandem cycling. One possible explanation is that the current protocol emphasised collaborative effort prior to warm-up, which may have reduced cadence variability between solo and tandem conditions.

### Pilot-stoker differences

Across all tandem configurations, a consistent asymmetry emerged between the two positions. Pilots maintained PO comparable to solo cycling, whereas stokers produced less PO relative to their solo performance. Despite this difference in PO contribution, heart rate and RPE did not differ significantly between pilots and stokers. Thus, the primary positional asymmetry was expressed in mechanical output rather than in average cardiovascular or perceptual load.

When compared to solo cycling, pilots exhibited lower heart rate while sustaining similar power output. As mechanical efficiency was not directly measured, this should not be interpreted as improved efficiency. Instead, the reduced heart rate may reflect differences in task regulation associated with the pilot position. Pilots are responsible for pacing and maintaining cadence and therefore function as the primary regulators of effort within the tandem. This regulatory role may allow for more anticipatory and progressive modulation of power output, potentially resulting in smoother effort distribution across the trial.

In contrast, stokers generated less power and reported lower RPE compared to their solo efforts. Because stokers do not control gearing, their contribution is inherently reactive to the pilot’s decisions. The lower RPE observed in stokers may therefore reflect differences in perceived responsibility or engagement associated with limited control [25-27], although reduced RPE is also consistent with their lower absolute power output. Importantly, the absence of heart rate differences between positions suggests that average physiological strain was comparable, even though the distribution of mechanical work differed.

Taken together, these findings indicate that position on the tandem shapes how effort is experienced and regulated within the tandem. Pilots function as proactive regulators of pacing within the tandem context, maintaining solo-level output while modulating cardiovascular strain relative to their solo condition. Stokers, in contrast, function as reactive contributors whose power output and perceived effort are influenced by the constraints of their position. This position-dependent pattern supports the view that tandem performance is not solely determined by individual capacity, but also by how responsibility and control are structured within the team.

### Pacing strategies

Pacing patterns were largely similar across conditions, with PO increasing in the last two minutes in all trials. The convergence of RPE across trials near the end of the effort suggests that, despite early perceived differences, riders in all conditions approached similar levels of perceived exertion as they neared exhaustion, consistent with the concept of centrally regulated pacing control [5, 28]. However, RPE was consistently lower in tandem conditions, particularly early in the trial, and may therefore reflect altered perception of effort due to shared workload [29, 30]. In addition, pilots appear to assume a more active role in the beginning of the time trial in unequal-capacity tandem conditions, which is even more pronounced when the pilot has a lower capacity than the stoker. However, in equal-capacity tandem conditions, pilots appear to have a more conservative start than the stokers, which may contribute to the lower mean power output in equal-capacity tandems when compared to solo. Although we can only speculate why pilots in equal-capacity tandem conditions started more conservative than during their solo trials, one plausible explanation is that evenly matched riders lack a clear performance hierarchy. This may introduce uncertainty about optimal pacing and lead to a more cautious start. Altering pacing patterns have been observed when athletes regulate effort under uncertainty or under influence of the presence of another athlete [31, 32]. With increased tandem experience, riders may learn to synchronise pacing more closely, reducing conservative starts and improving team performance [33].

### Practical Applications

From a coaching perspective, pairing cyclists of differing capacities may be acceptable for short-duration tandem efforts, as overall PO was not significantly affected by matching condition. However, coaches should monitor individual contributions within pairs, particularly stokers, to ensure balanced workload distribution. Providing real-time feedback on individual PO may help maintain engagement and align effort between riders [34]. Additionally, the conservative pacing seen early in tandem efforts suggests the need for practice sessions emphasizing start-phase power and intensity matching during acceleration.

### Limitations and future directions

This study involved trained but tandem-inexperienced cyclists performing short (10 minutes) laboratory time trials, which may limit generalisability to elite or Paralympic tandem teams in time trials (usually about 20 to 40 minutes) and road race conditions (1.5 to 2.5 hours), although the individual pursuit on the track falls within our set time frame. Communication between partners was not systematically recorded, and motivation levels were also not assessed, though they could substantially influence individual effort distribution.

As tandem cycling in a competitive setting is predominantly practiced within Paralympic sport, this study did not include stokers with a visual impairment, which limits the direct generalisability of our findings to Paralympic tandem teams. However, the controlled insights from our current sample provide a baseline for understanding position- and partner-dependent patterns that can inform hypotheses in Paralympic contexts. In the Paralympic context, pilot and stoker positions are fixed and not interchangeable, and stokers with a visual impairment usually also have limited access to visual feedback from the cycle computer display, depending on the degree of visual disability, potentially influencing pacing behaviour and perceived effort differently than in the present sample.

Including visually impaired stokers in the current study would have introduced additional practical constraints. Given the limited availability of experienced pilot–stoker combinations, this would likely have resulted in a substantially smaller sample size and insufficient statistical power. For these reasons, we chose to investigate the effects of partner ability and rider position in a controlled sample of trained but tandem-inexperienced cyclists.

Although caution is warranted when extrapolating these findings, the present results still offer relevant insights for Paralympic tandem cycling. Individuals with a visual impairment have been shown to rely more strongly on external guidance and following behaviour during coordinated movement tasks [35, 36], which may parallel the reduced power contribution observed in stokers in the current study. This suggests that the position-dependent differences identified here could be even more pronounced in visually impaired stokers under competitive tandem conditions.

Future research should directly examine whether similar patterns in PO, pacing, and perceived exertion emerge in Paralympic tandem teams, while also investigating the contribution of psychological and communication factors. Additionally, longer-duration and on-road tandem efforts involving elite teams are needed to verify the ecological validity of the present laboratory findings.

## Conclusion

Tandem performance emerges from the interaction between individual physical capacity, position on the tandem, and the regulation of shared effort between the riders. Although partner ability might be expected to influence performance, total PO was similar in equal and unequal pairings. However, stokers consistently contributed less and reported lower perceived effort, highlighting a systematic asymmetry in individual engagement within the tandem. These findings suggest that training strategies focusing on communication, pacing behaviour, and feedback on individual contribution may help optimise tandem performance. Although the present results were obtained in trained but tandem-inexperienced cyclists without visual impairment, they provide a controlled baseline for understanding how partner capacity and rider position shape tandem cycling performance, against which future studies in Paralympic tandem teams can be interpreted.

## Acknowledgements

The authors would like to thank the Royal Dutch Cycling Union (KNWU) for providing the road tandem bicycle.

## Supporting information captions

**Fig S1 A. Relative power over time for pilots and stokers in equal-capacity tandems**

**Fig S1 B. Relative power over time in higher-capacity pilots and stokers in unequal-capacity tandems**

**Fig S1 C. Relative power over time in lower-capacity pilots and stokers in unequal-capacity tandems**

